# Estimating Genetic Similarity Matrices using Phylogenies

**DOI:** 10.1101/2020.07.30.229286

**Authors:** Shijia Wang, Shufei Ge, Caroline Colijn, Priscila Biller, Liangliang Wang, Lloyd T Elliott

## Abstract

Genetic similarity is a measure of the genetic relatedness among individuals. The standard method for computing these matrices involves the inner product of observed genetic variants. Such an approach is inaccurate or impossible if genotypes are not available, or not densely sampled, or of poor quality (for example, genetic analysis of extinct species). We provide a new method for computing genetic similarities among individuals using phylogenetic trees. Our method can supplement (or stand in for) computations based on genotypes. We provide simulations suggesting that the genetic similarity matrices computed from trees are consistent with those computed from genotypes. With our methods, quantitative analysis on genetic traits and analysis of heritability and co-heritability can be conducted directly using genetic similarity matrices and so in the absence of genotype data, or under uncertainty in the phylogenetic tree. We use simulation studies to demonstrate the advantages of our method, and we provide applications to data.

## 1 Introduction

The computation of genetic similarities among samples (or taxa, or subjects, or individuals) is a key step in the analysis of quantitive genetic traits and in estimating the heritability of traits through variance component approaches (Chen and Witte, 2007; Malo et al., 2008; Tzeng and Zhang, 2007). For example, linear mixed models (LMMs) are popular methods for genomewide association studies, and matrix-variate normal methods are popular for heritability and co-heritability analyses. Both of these approaches demand efficient methods to determine genetic relatedness among samples (Kang et al., 2010; Lippert et al., 2011; Listgarten et al., 2013). The genetic relatedness among samples is usually specified through a genetic similarity matrix (Patterson et al., 2006; Thompson, 2013) derived empirically from genetic sequences, or from a kinship matrix (Boyce, 1983; Kirkpatrick et al., 2019) derived from a pedigree. Genetic sequences are often described by a series of genome locations at which mutations may be observed in the ancestry of the subjects. We consider single nucleotide polymorphisms (SNPs), locations at which a single DNA basepair can appear with more than one form (or, allele). The most common form is known as the major allele, and the less common forms are the minor alleles. We refer to the matrix resulting from the inner products of genetic sequences as the *empirical genetic similarity matrix*. This matrix is formed according to the following definitions. For the SNP at locus *m*, let *G_m_* denote the column vector of alleles. The genetic similarity between samples *i* and *j* for a haploid sample is defined as:

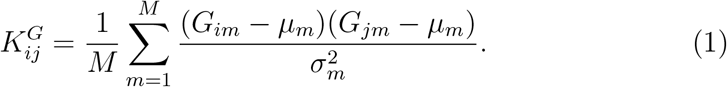

Here *G_im_* is the genotype of leaf *i* at marker *m* (*G_im_* ∈ {0, 1}), and *μ_m_*, 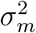 are the empirical mean and variance of the SNPs at marker *m*, and *M* is the total number of loci genotyped (Patterson et al., 2006). We will assume that *G_im_* = 1 indicates the event that sample *i* inherits the minor allele at marker *m*, and *G_im_* = 0 indicates the event that sample *i* inherits the major allele at marker *m*. The minor allele is defined as the allele that occurs less often among all of the samples. Note that permutations of the loci do not affect the empirical genetic similarity. This formulation of genetic similarity is used in linear mixed models (Listgarten et al., 2012), and multi-phenotype models (Yang et al., 2011). The matrix entry 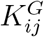 represents the expected genetic contribution to the correlation between phenotypes for samples *i* and *j*, and thus *K^G^* may be used as a fixed effect. Other methods for computing relatedness (for example patristic distance or identity by descent) have closed forms, but are not directly exploitable in fixed effects models.

The computation of genetic similarity among samples using equation (1) requires genotyped sequences, and several challenges may arise in the computation. Firstly, the genetic sequences may not be readily available. This may be the case when extinct species are examined. Secondly, it is hard to assess the uncertainty for empirical genetic similarities in cases for which the genotyped sequences have low quality, or are homoplastic. This case may arise when examining bacterial genomes or *de novo* sequences. These challenges motivate us to propose an approach that does not involve genetic sequences.

In this article, we develop a method for estimating expected values for the entries of a genetic similarity matrix using a phylogeny (*i.e*., a closed form for the expected value of equation 1, conditioned on a tree). Our proposed approach does not require that the sequences for individuals be genotyped (or measured). Our method allows analysis based on equation 1, such as analysis using fixed effects models, to proceed for situations in which genotypes are not available but an approximation of the molecular phylogeny is, or for situations in which there is uncertainty in equation 1 stemming from low quality genotyping or short genomes. For example, in some studies of extinct species, genotypes might not be available but an approximate molecular phylogeny, based on morphological data and geological dates, might be. Instead of requiring individual genotypes, we assume that a phylogeny is given.

The relatedness among samples is computed by integrating over all mutations occuring in the branches of a tree, under the assumption of an infinite sites model (Ma et al., 2008), with a constant mutation rate for the evolutionary process. The infinite sites model is a popular and simple model for mutations in genome evolution, and it is a reasonable model when the genome is large. The infinite sites model postulates that new mutations are always at novel loci (and not re-entrant). Genetic recombination among samples is not considered in this approach (however, the approach for samples with recombination is clear), as we mainly consider genetic similarities at the species level. We numerically demonstrate that our proposed *expected genetic similarity matrix* is asymptotically equivalent to the empirical genetic similarity matrix, with an infinite number of genotyped loci (provided that the tree is correct).

The main application of our work is to provide a correspondence between phylogenetic trees and the normalized genetic similarity matrices (Patterson et al., 2006) that are standard in linear mixed models and heritability analysis. Often, many sampled molecular phylogenies are provided conditioned on genotypes (for example with BEAST: the Bayesian Evolutionary Analysis Sampling Trees software; Drummond and Rambaut 2007). Our work allows computation of the expected value of the standard and normalized genetic similarity matrix (Patterson et al., 2006) for each of the sampled trees. These matrices can then be used to average over linear mixed models applied to each sample (respecting the uncertainty in the genetic similarity matrix), or they can be used in Bayesian models extending the usual linear mixed model paradigm. In contrast, work based on computing equation 1 directly from genotypes provides a point estimate of the genetic similarity matrix (without respecting the uncertainty implied by the genotypes).

In our simulations, the expected genetic similarity matrix is invariant to the number of samples and to the mutation rate in the infinite sites models. Our approach is more accurate than the multidimensional scaling (MDS) approach implemented in the software package *pyseer* (Lees et al., 2018) and Gaussian distance similarity matrices (González-Recio et al., 2008; Ickstadt et al., 2005). The MDS approach for genetic similarity matrix estimation also operates on phylogenies, but is not based on integration over all mutations.

We use our approach to compute the genetic similarity matrix for a set of ancient hominin species, with the phylogeny found using BEAST on the set of morphological phenotypes used in Dembo et al. (2016). Genetic similarity matrices are often used to partition the covariance matrix into additive components that arise from genetic, environmental or random factors (Dahl et al., 2016; Wang et al., 2011). Given this estimated genetic similarity matrix, we compute the component of genetic covariance that is inherited along the tree (before and after speciation events) for the heights of the ancient hominin species. In particular, we apply a linear mixed model (Yang et al., 2011) to compartmentalize the covariance matrix of the heights of the species, such that the genetic component is a scaled version of our estimate for the genetic similarity matrix. This demonstrates how our estimate can be used to determine genetic aspects without access to measured genotypes (instead, with access to a tree which we assume approximates the correct phylogeny). In addition, we apply our algorithm to evaluate the uncertainty of genetic similarities among hominin species.

### 1.1 Related work

Genetic similarity is a basic aspect of many approaches in genetic analysis. Identity-by-descent (IBD; Whittemore and Halpern 1994) is a popular measure for genetic similarity, based on identification of stretches of genetic sequences which have identical ancestral origin. Kinship coefficients describe the probability that two random alleles from a pair of individuals are IBD, and these coefficients are commonly used to measure genetic similarity between a pair of individuals. There are several ways to compute kinship coefficients, including from genetic data, and also based on pedigree graphs Maruyama and Yasuda (1970). The idea of using trees to define similarity matrices (for use as fixed effects) is explored in Housworth et al. (2004). But that work uses estimates of similarity matrices based on pedigrees (and does not use the definition of genetic similarity as a normalized inner product of genetic sequences, as is standard in applications of LMMs to genome-wide association studies; Listgarten et al. 2013). Similarly, in Abney (2009) a novel graphical algorithm for computing the kinship coefficients using graph traversal is developed: “kinship graph”, and in Thornton et al. (2012) a robust method “REAP” is developed, to estimate IBD-sharing probabilities and kinship coefficients for admixtures. However, neither of these methods are designed to estimate the normalized inner product of genetic sequences (Equation 1).

Computational concerns for kinship matrices have also been considered: Kirkpatrick et al. (2019) propose a fast algorithm for calculation of kinship coefficients for individuals in large pedigrees. Our work provides an algorithm with similar computational complexity.

In further lines of research, if genetic sequences are observed, a variety of measures have been used to compute genetic similarities using categorical data clustering (González-Recio et al., 2008; Ickstadt et al., 2005) by creating dissimilarity scores for genotypes and then converting the dissimilarity scores to similarity scores. For example, Murray et al. (2017) present the k-mer Weighted Inner Product (kWIP), an assembly based estimator of computation of genetic similarities among individuals, that is also alignment-free. The pairwise similarity is obtained from k-mer counts. In contrast, less work has been done in assessing the statistical relationship between phylogenetic trees (rather than pedigrees or alignment-free methods) and genetic similarity matrices. The most similar approach is based on MDS, which has been applied to the control of linear mixed models (Lees et al., 2018). This MDS approach also provides a deterministic function that outputs a similarity matrix given a fixed tree. In typical protocols, the fixed tree used in the production of the similarity matrix may be produced from genotypes using BEAST.

## 2 Methods

Let *T* be an unrooted binary tree with leaves *i* ∈ {1, …, *N*}. The phylogeny *T* represents relationships among *N* taxa through a tree topology *τ* and a set of branch lengths ***e*** = (*e*_1_, *e*_2_, …, *e*_2*N*–3_). The leaves of a sampled tree *T* are the samples in the study. Each interior node of *T* represents the most recent common ancestor of the two children of that node, and the branch lengths are proportional to the evolutionary distances between pairs of nodes. An unrooted tree represents the relatedness of leaves without making assumptions about an earliest common ancestor. In contrast, a rooted binary tree describes the relatedness for a set of leaves in the tree from a single common ancestor at the root.

In computational genetics, the infinite sites model is commonly used to model genetic variation (Kimura, 1969). In this model, we assume polymorphism arises by single mutations of unique sites at locations within the genetic sequence, with all mutations occurring at different positions, implying that all genetic variants are biallelic. Let **G** = [*G*_1_, *G*_2_, …, *G_M_*] denote genotype data observed at *M* genetic variants, with *G_m_* denoting a column vector of alleles for the *m*-th SNP for all *N* subjects. The genetic similarity matrix of the leaves (Patterson et al., 2006) is an *N* × *N* symmetric matrix *K* defined in equation (1), in which 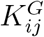 denotes the genetic similarity between sample *i* and sample *j*. Our goal is to compute the expected value of 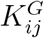 given a tree *T*. By additivity of expectations, from equation (1) we arrive at the following expectation through integration over all mutations:

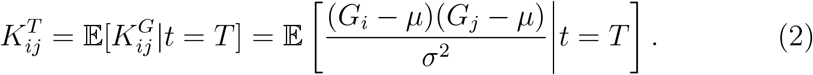

Here *G_i_* is a random variable giving the genotype of a marker placed at a random location of the tree, and *μ*, *σ*^2^ are random variables giving the mean and variance of *G_i_* as an element of the set {0, 1} (assuming haplotype data).

To compute equation (2), we integrate the location of the marker over the tree, noting that expectation splits linearly over the union of the domain of integration. Assuming a neutral model with a constant mutation rate, the values of *G_i_* are completely determined by the location of the marker, and they are constant over each edge of the tree. So, equation (2) can be rewritten as a weighted sum over edges of the tree. The weights are given by |*e_d_*|/|*T*|, where |*e_d_*| is the branch length of an edge *e_d_*, and |*T*| is the sum of the branch lengths of all edges in the tree. The expected value on the right hand side of equation (2) is thus given by the following:

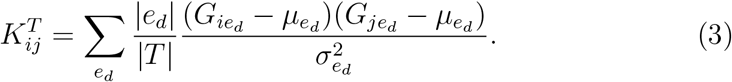

The values *G_ie_d__*, *μ_e_d__*, 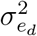 are found by considering a mutation on each edge *e_d_*. Note that Equation (3) is an approximation of expected value of genetic similarity found by assuming that the tree provides the correct phylogeny. Hence, it represents the genetic similarity between taxa *i* and *j*. Under our assumptions, *μ_e_d__*, 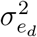 are summary statistics for *G*, and *K^T^* is a deterministic function of these variables according to equation (1). These values can be computed by considering the following steps for each element of the sum from equation (3).

1. Let *c*_0_ and *c*_1_ be the bipartition of taxa induced by segregation over edge *e_d_* in the unrooted tree *T*. Figure 1 shows an example of this operation. Without loss of generality, we assume that the number of leaves in *c*_0_ is greater than or equal to that of *c*_1_. This means that each leaf in *c*_0_ will have the major allele, and each leaf in *c*_1_ will have the minor allele.
2. The genotype *G_ie_d__* is 1 if leaf *i* is in *c*_1_ and *G_ie_d__* is 0 if leaf *i* is in *c*_0_.
3. The mean *μ_e_d__* is the number of leaves in *c*_1_, divided by the total number of leaves.
4. 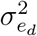 is the variance of the Bernoulli distribution implied by the allele:

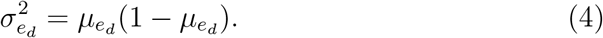

**Figure 1:**
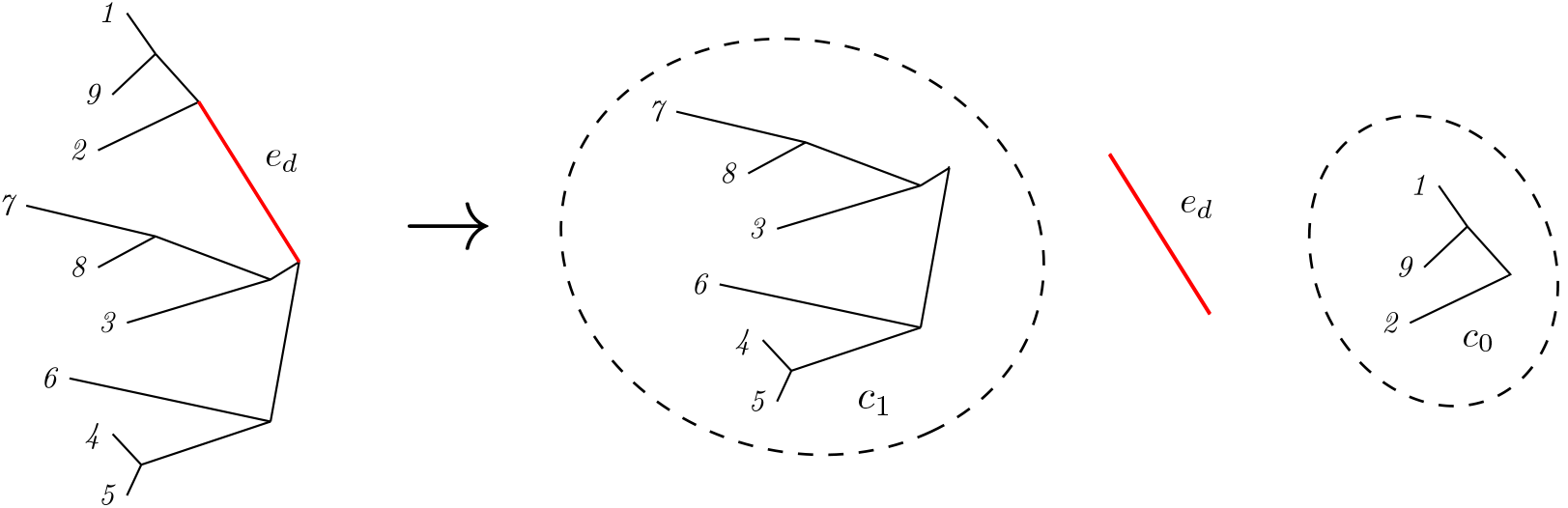
Bipartition of taxa induced by an unobserved genetic variant on edge *e_d_*. Only the membership of each taxon in the bi-partitioned set is needed in the computation of the expected mean *μ_e_d__*, and variance 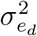.

Note that if *c*_0_ and *c*_1_ are the same size, then *μ_e_d__* = 1 – *μ_e_d__* = 0.5, and 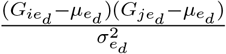 does not depend on which allele is assigned to 0 or 1 (*i.e*., breaking ties for *c*_0_ and *c*_1_ one way or the other leaves 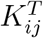 unchanged). Equation (3) thus suggests the following 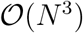 algorithm (Algorithm 1) for computing the expected genetic similarity matrix, given the tree *T*. In this algorithm, *A*’ denotes the transpose of the matrix *A*. We provide an open source software implementation for this method^1^.

**Algorithm 1.**
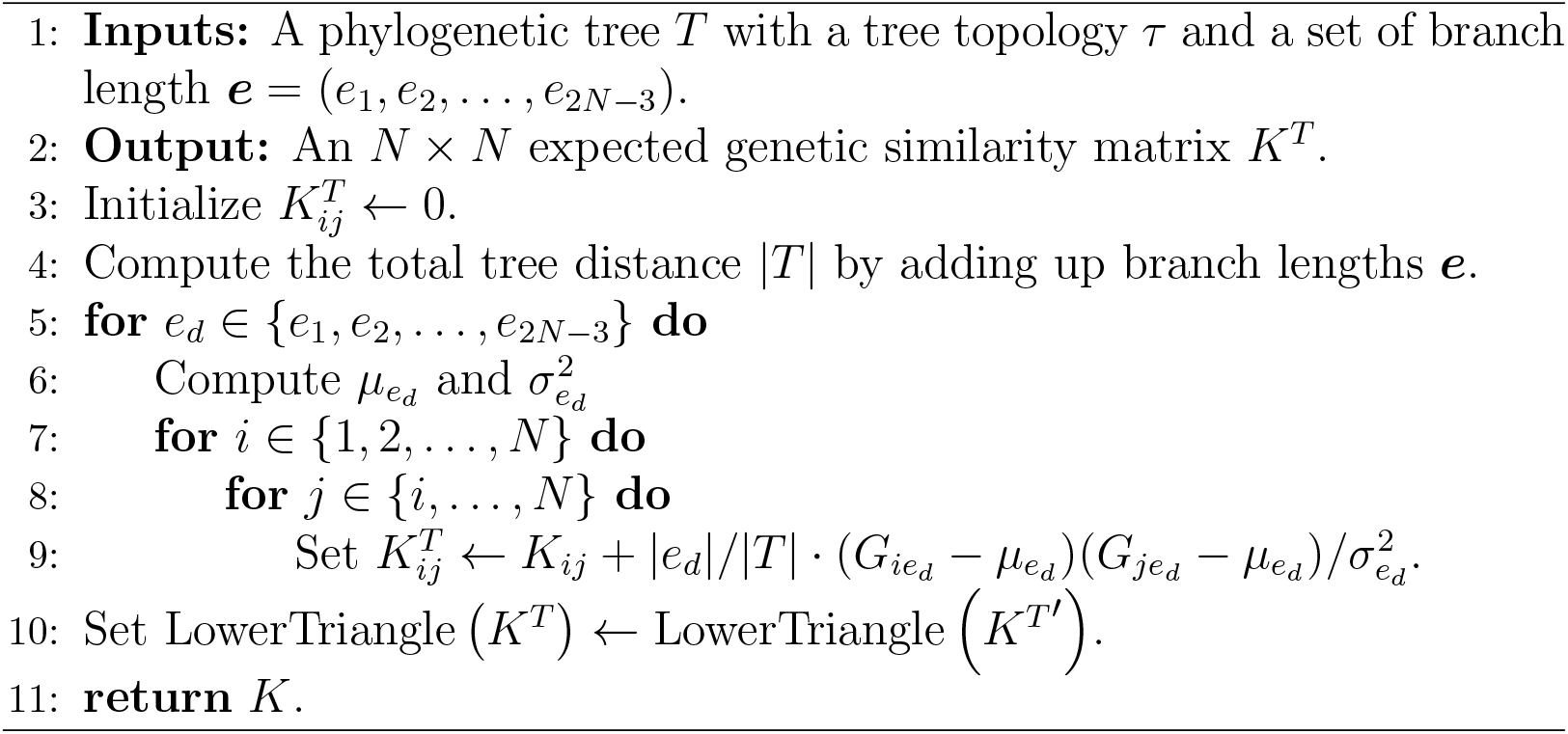
From phylogeny to expected genetic similarity matrix

## 3 Experiments

In our numerical experiments, we use the *ms* software (Hudson, 2002) to simulate binary trees and genetic variation (*ms* is a coalescent simulator that generates genetic samples under the assumption of the neutral model and the infinite sites assumption). We also assume constant effective population size *N*. The branch lengths are thus in units of 2*N* generations. We simulate genetic sequences for haploidy of population and assume no recombination during the history (this assumption is appropriate for evolutionary timescales with species for which lateral transfer is negligible).

### 3.1 Simulation 1

In this simulation study, we numerically demonstrate the consistency of our algorithm. We simulate datasets for three scenarios: A, B, C. In scenario A, we simulate 100 trees, with *N* = 20 samples (or subjects, or taxa) for each tree. Also, for each tree, we simulate two sets of genetic sequences, each with *M* = 1000 or 20 loci. In scenario B, we simulate 100 trees with *N* = 6 taxa, each with *M* = 1000 loci. In scenario C, we simulate 100 trees with *N* = 20, *M* = 1000, and for each tree we simulate SNPs with a neutral mutation rate of *μ* = 7.5 mutations on the entire sequence per generation. In the *ms* software, the mutation parameter *θ* = 2*Nμ* (Hudson, 2002).

For each scenario, we compute the empirical genetic similarity matrix using the simulated genotypes to form a ground truth. Then, we compute the expected genetic similarity matrix using Algorithm 1 and the simulated tree. Figure 2 displays scatter plots comparing our method for computing genetic similarity matrices to the ground truth (the ground truth is found by applying equation (1) to the simulated genotypes). Figure 2 shows that for a higher value of *M*, the correlation among the entries of the genetic similarity matrices computed from trees and from genotypes may be stronger than it is for low values of *M* (the number of loci). This figure also shows that even a small number of samples may produce unbiased estimates of the genetic similarity matrix. We also show that for situations with small numbers of samples, the estimates may still improve for higher values of *M*.

**Figure 2:**
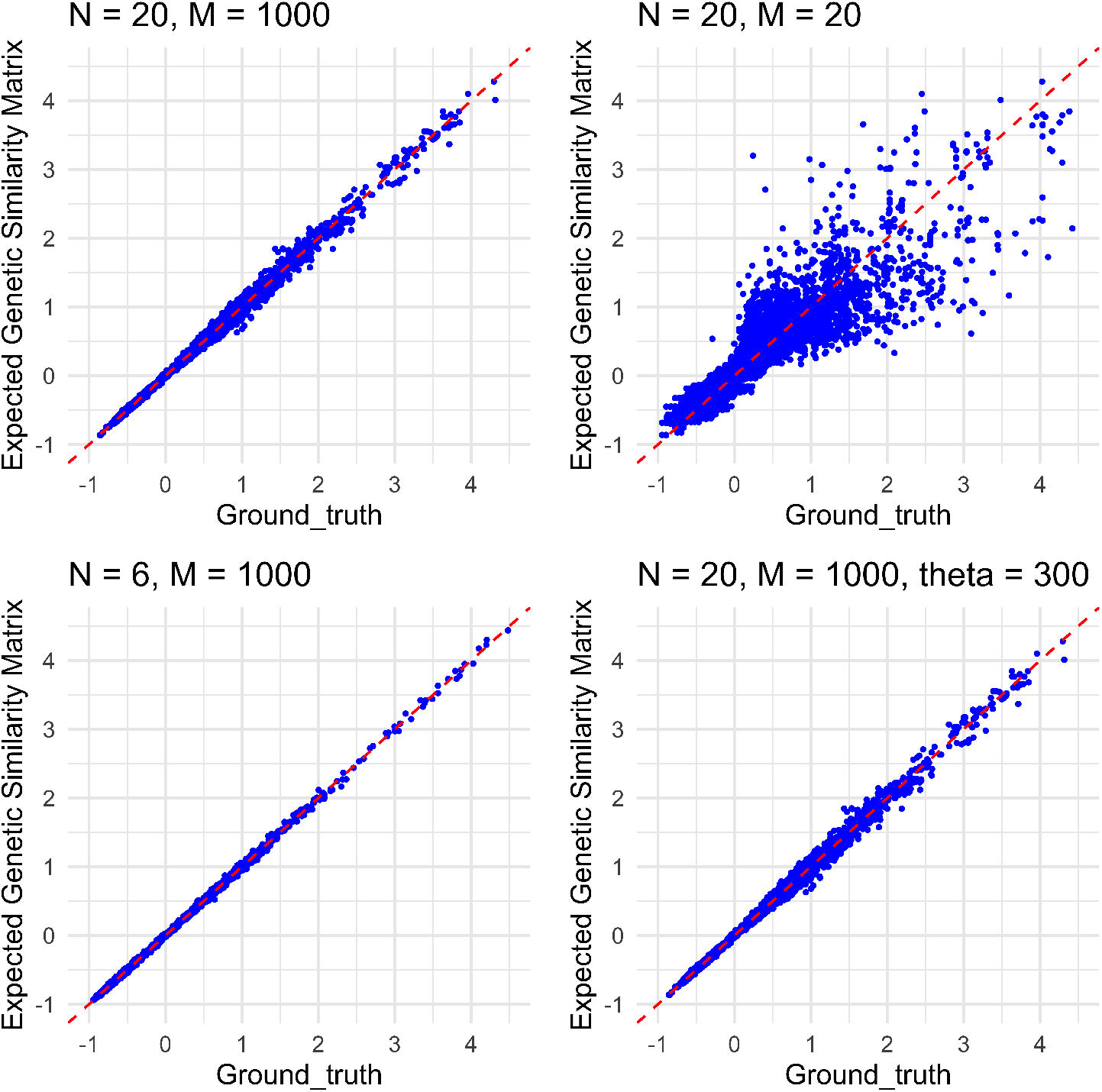
Comparison of simulated genetic similarity matrices from trees and genotypes. Scatter plots are provided for entrywise differences between the genetic similarity matrices produced by our method and the ground truth (from equation 1, applied to the simulated genotypes). The *top left* and *top right* panels show the entrywise differences for the two conditions for scenario A, and the *bottom left* and *bottom right* panels show the entrywise differences for scenarios B and C (respectively).

The entries of genetic similarity matrices computed from trees and genotypes become closer together as we increase *M* on the genotype from 20 to 1000, providing evidence for consistency of our methods over the range of *M* considered. In addition, these simulations suggest that the expected genetic similarity matrix is invariant to the number of individuals and the neutral mutation rate.

We conduct an additional set of experiments to investigate the difference between entries of the expected genetic similarity matrix and the ground truth as (*A*): a function of number of loci for each sequence; (*B*): a function of number of samples (or subjects, or taxa). For (*A*), we simulate 100 trees, with *N* = 50 samples (or subjects, or taxa) for each tree. In addition, we simulate eight sets of genetic sequences, each with *M* = 10, 30, 100, 300, 1000, 3000, 10000, or 30000 loci, for each tree. Figure 3 displays the difference between the expected genetic similarity matrices and the ground truth with different numbers of loci for each fixed number of sequences. The violin plots denote the entrywise difference between the expected genetic similarity matrices and the ground truth 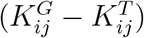. This figure indicates that for a higher value of *M*, the correlation among the entries of the genetic similarity matrices computed from trees and from genotypes is larger than it is for low values of *M*.

**Figure 3:**
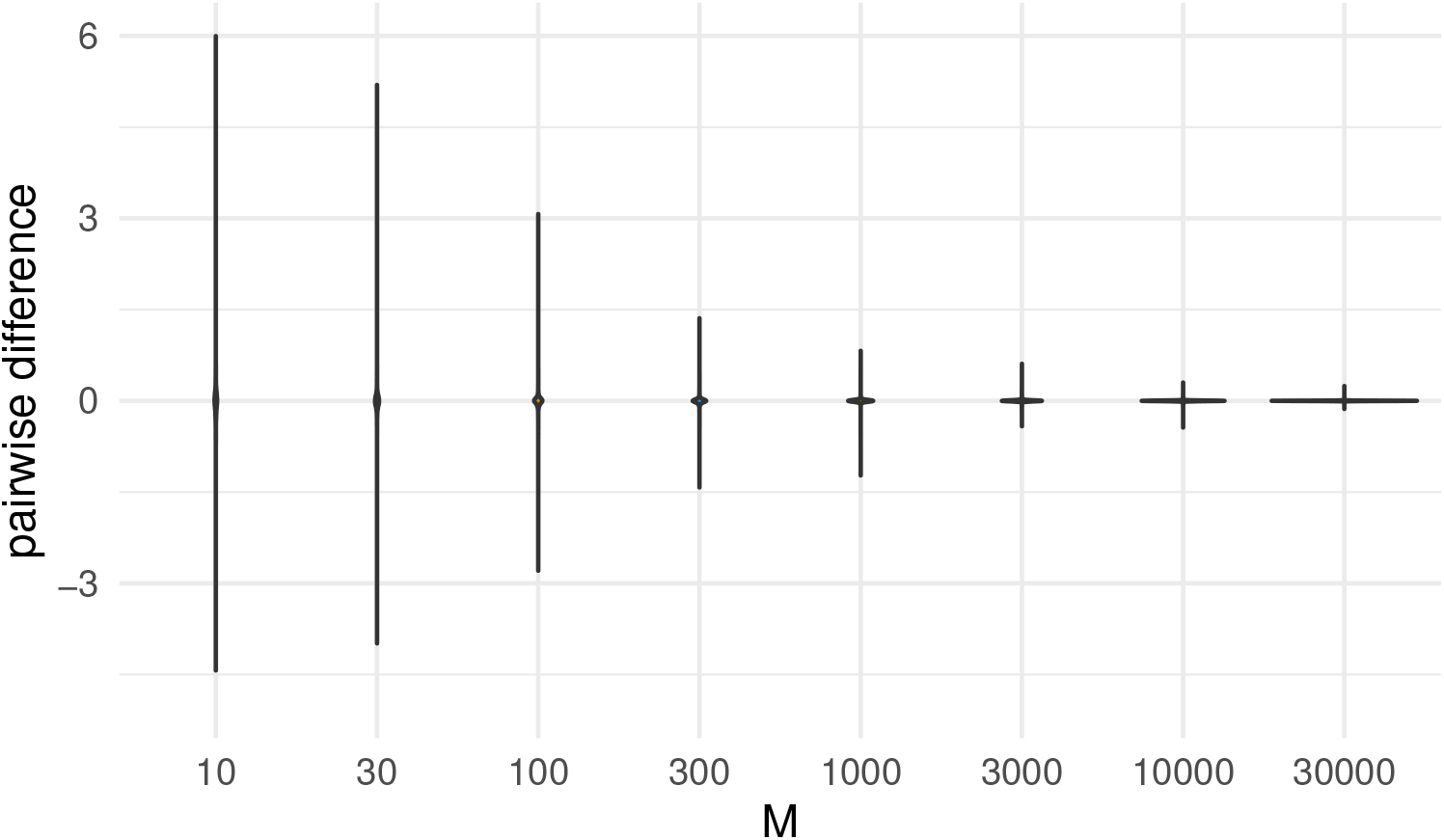
Comparison of simulated genetic similarity matrices from trees and genotypes. The violins are provided for entrywise differences between the genetic similarity matrices produced by our method and the ground truth.

For (*B*), we simulate six sets of trees, each with *N* = 10, 30, 100, 300, 1000, 3000 taxa for each tree, and with 100 trees for each set. We simulate genetic sequences, with 1000 loci, for each tree. Table 1 shows the summary statistics (mean, standard deviation, 2.5%, 25%, 50%, 75%, 97.5% quantiles) for entrywise differences between the genetic similarity matrices produced by our method and the ground truth 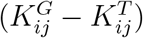. This table indicates that the correlation among the entries of the genetic similarity matrices computed from trees and from genotypes is invariant to the number of taxa (*N*). This invariance is further supported in Figure S1 of the Supplementary Material through identical violin plots.

**Table 1:** Comparison of simulated genetic similarity matrices from trees and genotypes as a function of number of taxa. The summary statistics (mean, standard deviation, 2.5%, 25%, 50%, 75%, 97.5% quantiles) for entrywise differences between the genetic similarity matrices produced by our method and the ground truth are close. This observation is further supported in Appendix B of the Supplementary Material.

| #taxa | mean | SD | 2.5% | 25% | 50% | 75% | 97.5% |
| --- | --- | --- | --- | --- | --- | --- | --- |
| 10 | 0.000 | 0.028 | -0.054 | -0.013 | 0.000 | 0.012 | 0.057 |
| 30 | 0.000 | 0.033 | -0.055 | -0.011 | 0.000 | 0.012 | 0.054 |
| 100 | 0.000 | 0.034 | -0.049 | -0.011 | 0.000 | 0.011 | 0.050 |
| 300 | 0.000 | 0.035 | -0.039 | -0.010 | 0.000 | 0.010 | 0.044 |
| 1000 | 0.000 | 0.034 | -0.041 | -0.009 | 0.000 | 0.010 | 0.036 |
| 3000 | 0.000 | 0.036 | -0.055 | -0.008 | 0.002 | 0.012 | 0.038 |

### 3.2 Simulation 2

In our second simulation study, we compare our tree based genetic similarity matrix approach to other approaches that measure genetic similarities using phylogenies. The first approach we compare to is a Gaussian distance similarity matrix (González-Recio et al., 2008; Ickstadt et al., 2005), in which we first compute the distance between two samples *i* and *j*, *d_ij_*, by computing the length of the shortest path between them on the tree. Then we convert the distance to a similarity matrix through the equation 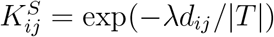. Here λ is a fixed bandwidth. We refer to Appendix A of the Supplementary Material for more details of this approach. This approach is similar to Ickstadt et al. (2005) and González-Recio et al. (2008), but with distances computed through the phylogeny. The second approach we compare to is the multidimensional scaling (MDS) approach implemented in the software package *pyseer* (Lees et al., 2018). In the MDS approach, the similarity between each pair of samples is calculated based on the shared branch length between the pair’s most recent common ancestor and the root. Multidimensional scaling is then performed on the resulting similarity matrix. We denote the genetic similarity matrix computed via *pyseer* by 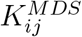 (note that we do not perform the MDS itself, and instead compare the similarity matrices directly).

We simulate 100 trees, with N = 20 taxa in each tree. For each tree, we simulate genetic sequences with *M* = 1000 loci. We compute the genetic similarities via 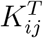 (our method, based on the tree), 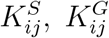 (the inner product from the sampled genotypes) and 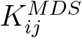. Figure 4 displays the comparison of genetic similarity matrices using trees. The violins denote the differences between 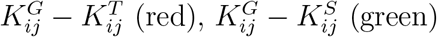 and 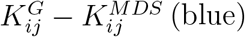. The empirical and expected genetic similarity matrices are closer to each other than they are to the Gaussian distance similarity matrix, and are closer to each other than they are to the MDS similarity matrix.

**Figure 4:**
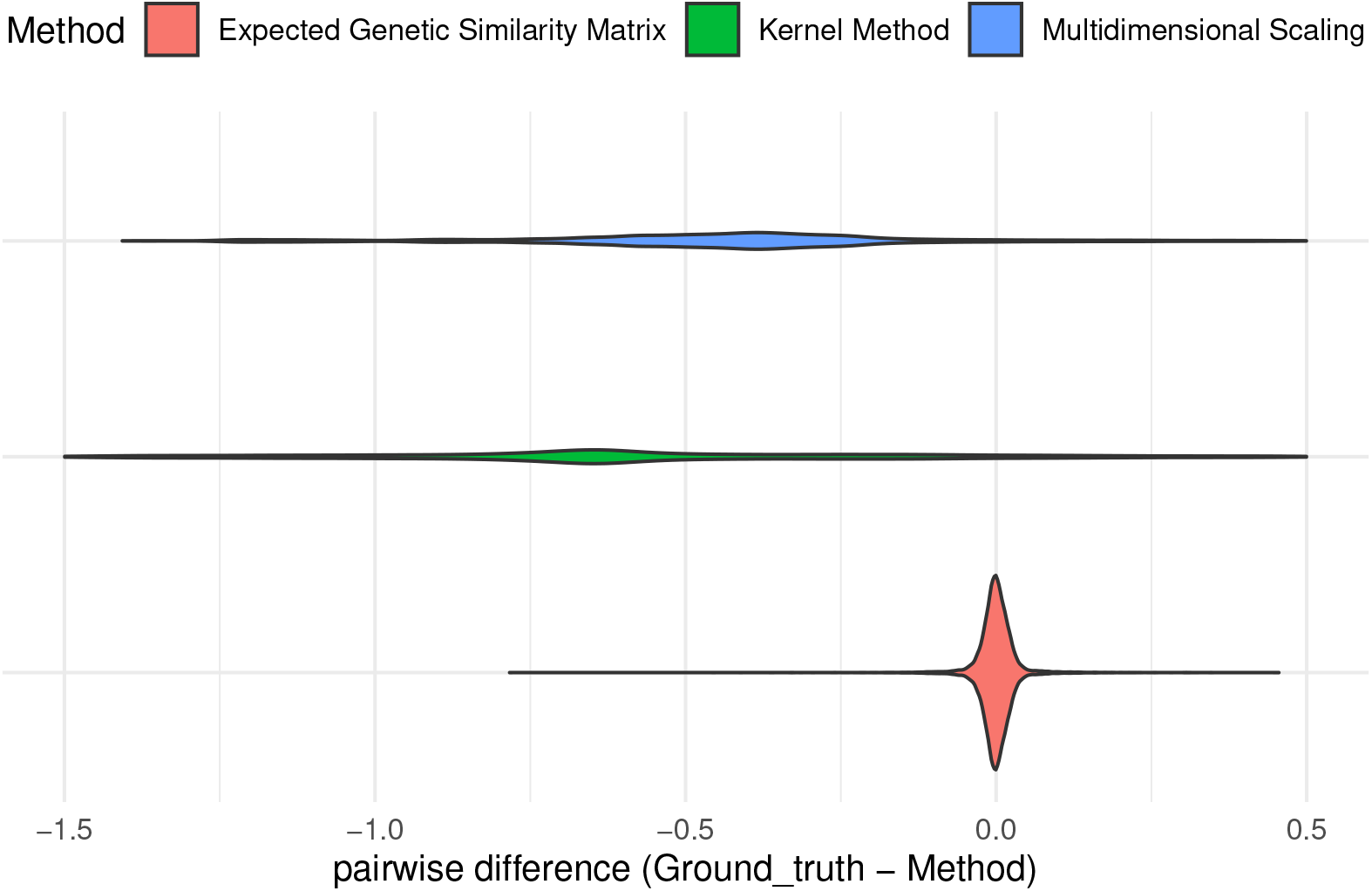
Comparison of expected genetic similarity matrix approaches: 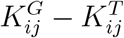 (our method, *bottom*), 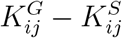 (*middle*) and 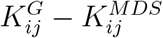 (*top*). The entries of the expected genetic similarity matrices (the red violin, thin) are close to the empirical genetic similarity matrix. They are closer to the empirical genetic similarity matrix than are the entries of the Gaussian distance similarity matrices (the green violin), or the multidimensional scaling similarity matrices (the blue violin).

Note that the simulation conditions of the comparison between our method and the empirical genetic similarity matrix are the same as the upper left panel of Figure 2. For the 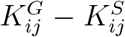 condition, we note that the median difference is far from zero (as the range of the Gaussian distance method is positive). However, 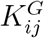 and 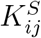 are still correlated (*ρ* = 0.27, *p* < 0.001). Here ρ denotes the correlation between entries of 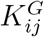 and 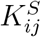, and *p* denotes the p-value for the null hypothesis that *ρ* is equal to 0. For the 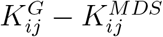 condition, the median difference is still far from zero, but less far than it is for the 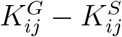 condition. The 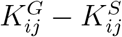 still involves correlation between the entries of the similarity matrices (*ρ* = 0.56, *p* < 0.001).

### 3.3 Ancient hominin data

We consider two experiments on ancient hominin data. In the first experiment, we apply our method to compute the genetic similarities between 8 hominin species. Fossil remains of a previously-unknown ancient hominin species (*Homo naledi*) were discovered in South Africa (Berger et al., 2015) and the DNA of this ancient hominin species remains unsequenced. While some ancient hominin species have been sequenced, often geological, paleontological, morphological or anthropological observations are used to assess the evolutionary relationships among extinct species (Dembo et al., 2016). In this way, a tree *approximating* the molecular phylogeny can still be constructed. We assume that the branch lengths of the molecular phylogeny are proportional to the phylogeny inferred from paleontological dates (and in experiments described later in this section we assume that molecular phylogeny is proportional to morphological trees). Recent work in hominin phylogeny suggests that morphological trees broadly match molecular phylogeny (Wiens, 2004; Wood and Boyle, 2017). However, convergent evolution and high substitution rates modulate the validity of this assumption (Berger et al., 2017; Scally et al., 2012). Our method allows us to compute genetic similarity matrices which could then be used in analysis of heritability or coheritability (as is done in Dahl et al. 2016), in absence of any genetic sequences.

We use our proposed approach to compute the expected genetic similarity matrix for *H. sapiens* and 7 extinct cousins or ancestors of humans (H. *habilis, H. rudolfensis, Georgian H. erectus, African H. erectus s.s., Asian H. erectus s.s., H. antecessor, H. neanderthalensis*). The phylogenetic tree for the species that we consider are provided and computed in Dembo et al. (2016) using geological dates. While Dembo et al. (2016) reconstruct the phylogenetic tree of 24 species, we consider the 7 hominin species most closely related to humans, as well as humans themselves (yielding 8 species). Figure 5 shows the heatmap for the expected genetic similarity matrix for the 8 hominins considered in this article. The genetic similarities between *H. rudolfensis* and *Georgian H. erectus, H. rudolfensis* and *H. habilis, Georgian H. erectus* and *H. habilis* are larger than the rest.

**Figure 5:**
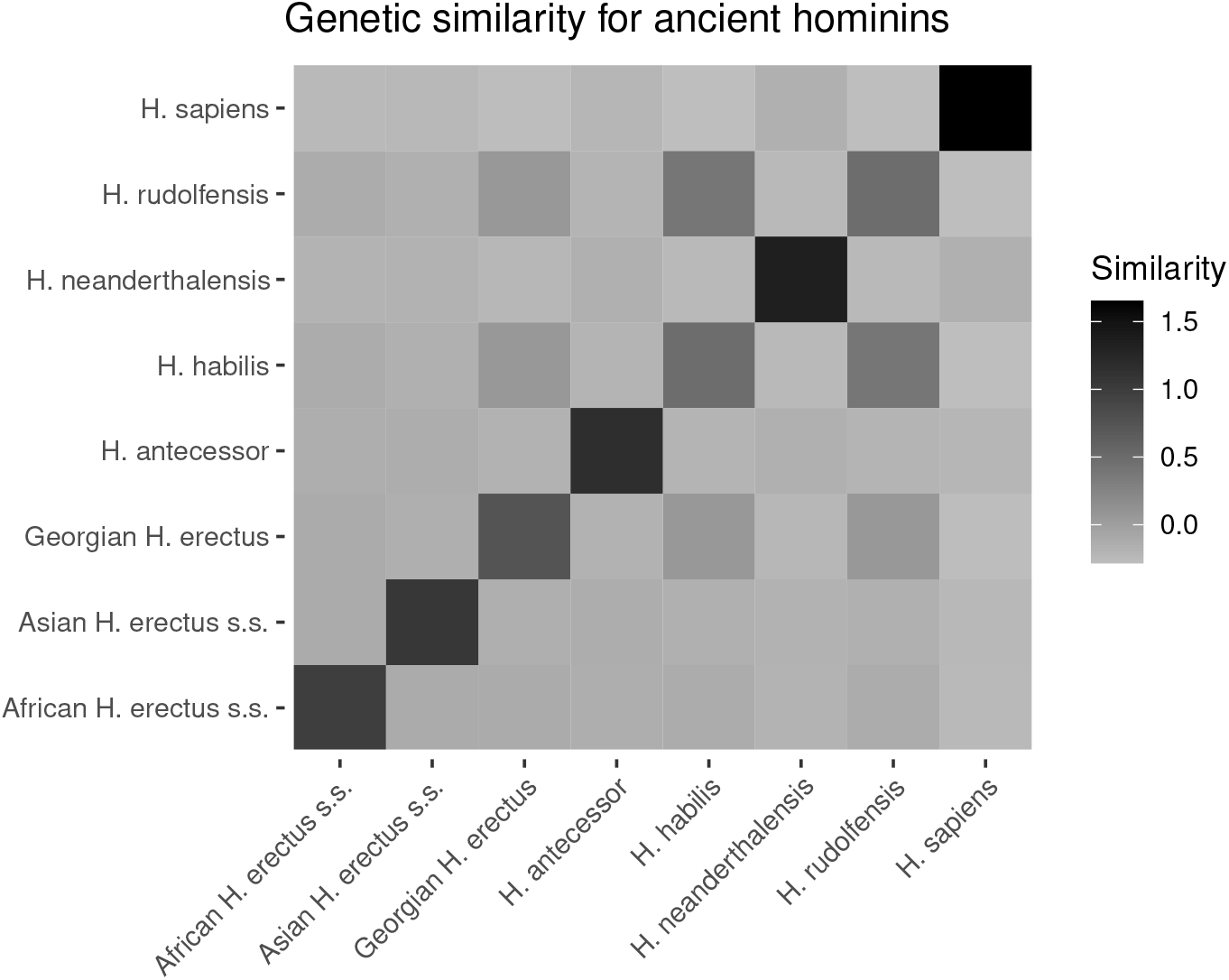
Genetic similarity matrix for 8 hominin species as a heat map, computed using geological dates and Algorithm 1.

We also provide the specific values of the heatmap in Figure 5 of the similarities in Table S1 of Appendix C of the Supplementary Material. This table could be used in conjunction with models such as Dahl et al. (2016) to conduct heritability or co-heritability analysis on traits of the species considered. To that end, we consider a heritability analysis on the average heights of genetic males in the 8 hominin species. We gather these average heights of ancient hominins from the following references: Carretero et al. (1999); Helmuth (1998); Lordkipanidze et al. (2007); McHenry and Coffing (2000). We estimate variance components from a linear mixed model (a standard model for methods such as genome-wide complex trait analysis or GCTA; Yang et al. 2011). The form of this model is as follows:

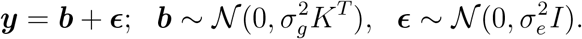

Here ***y*** denotes the standardized phenotypes (genetic male heights), and *I* is an identity matrix. The random effects ***b*** are the genetic effects. The term ***ϵ*** encodes the environmental effect.

Narrow sense heritability, denoted *h*^2^, is the fraction of the variance of *y* due to the genetic component. This quantity is defined as follows:

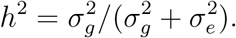

We fit the linear mixed model with *K^T^* (our expected genetic similarity matrix) computed earlier in this Section. We conducted inference using MCMC (Markov chain Monte Carlo; Gibbs sampling) for the parameters of the linear mixed model, using 10, 000 iterations, of which the first 5, 000 are discarded for burnin. Figure 6 shows the MCMC trace plots of 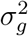 and 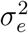. The red dashed lines indicate posterior means for 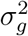 and 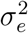 with the burnin discarded. Figure 7 shows the histograms for posterior samples of 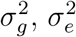 and *h*^2^ with the burnin discarded. The red lines indicate the posterior means, and the blue dash lines indicate the 95% credible intervals.

**Figure 6:**
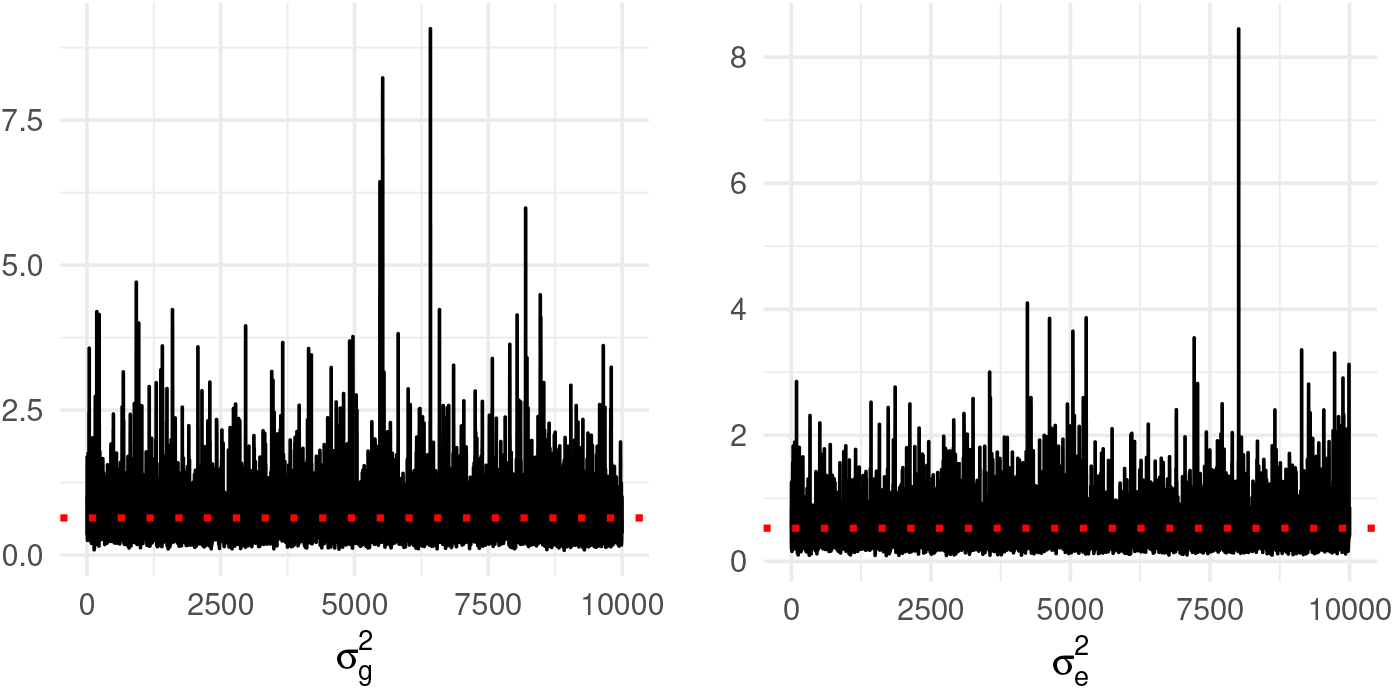
MCMC trace plots of 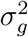 and 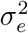. The red dashed lines indicate posterior means for 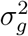 and 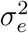 with initial 5, 000 iterations as burnin.

**Figure 7:**
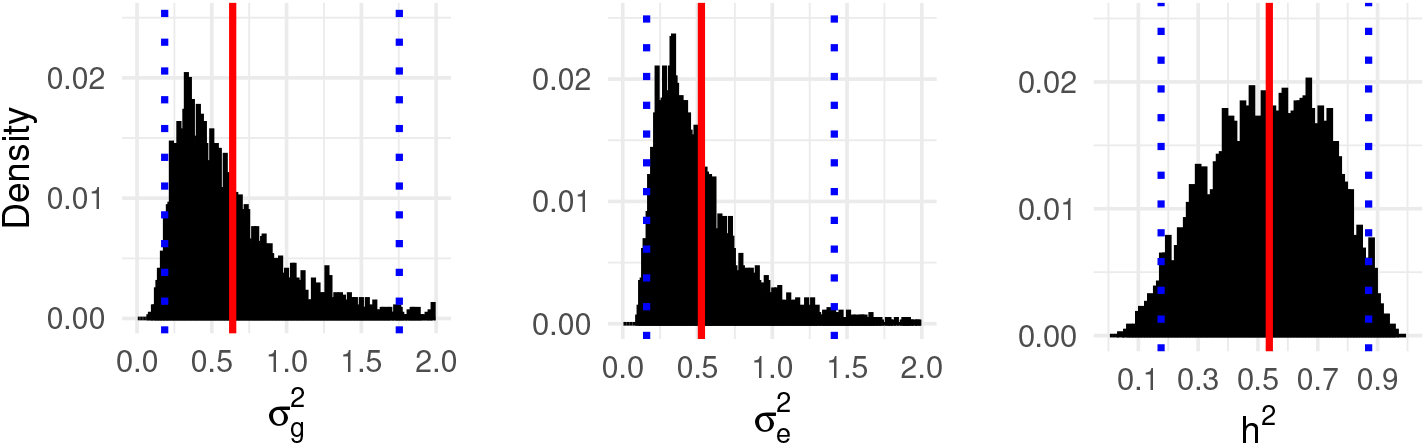
Histograms for posterior samples of 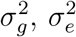 and *h*^2^. The red lines indicate posterior means for 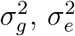 and *h*^2^, the blue dash lines indicate the 95% credible intervals.

We obtain posterior means and 95% credible intervals 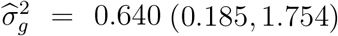 and 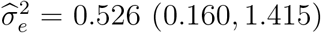. Therefore, from 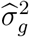 and 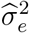 we estimate that the narrow sense heritability for this trait is 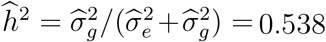, with 95% credible interval given by (0.176, 0.870). Note that the validity of these estimates depends crucially on the assumptions of the model: Firstly, that the paleontological tree is proportional to the phylogeny, and also that the assumptions of the neutral model and infinite sites model hold. For humans, the narrow-sense heritability of height suggested by Yang et al. (2015) is between 0.6 and 0.7.

### 3.4 Uncertainty assessment in hominin species

In this section, we present a second experiment on hominin data. We apply our approach to evaluate the uncertainty for genetic similarities for 24 hominin species. The DNA sequences of some hominin species remain unavailable (e.g. *Sahelanthropus tchadensis*). We compute the genetic similarity matrix for a posterior sample of phylogenetic trees for the hominin species. We used 10, 000 posterior tree samples (after thinning) created through MCMC with BEAST2 (Bouckaert et al., 2014). Details of the BEAST2 run are provided in Appendix D of the Supplementary Material. The morphological data used are provided in Dembo et al. (2016), but BEAST2 was used rather than Mr. Bayes 3.2.4 (Ronquist et al., 2012).

The resulting matrices represent the uncertainty of genetic similarities among the hominin species. Figure 8 displays the uncertainty of genetic similarities between *Ar. ramidus* and *Au. anamensis, G. gorilla* and *P. troglodytes, P. robustus* and *H. naledi, P. troglodytes* and *Au. afarensis*. We refer readers to Appendix E of the Supplementary Material (Figures S2-S4) for the posterior mean and the 95% credible interval for a heatmap of the genetic similarity matrix for all 24 hominin species.

**Figure 8:**
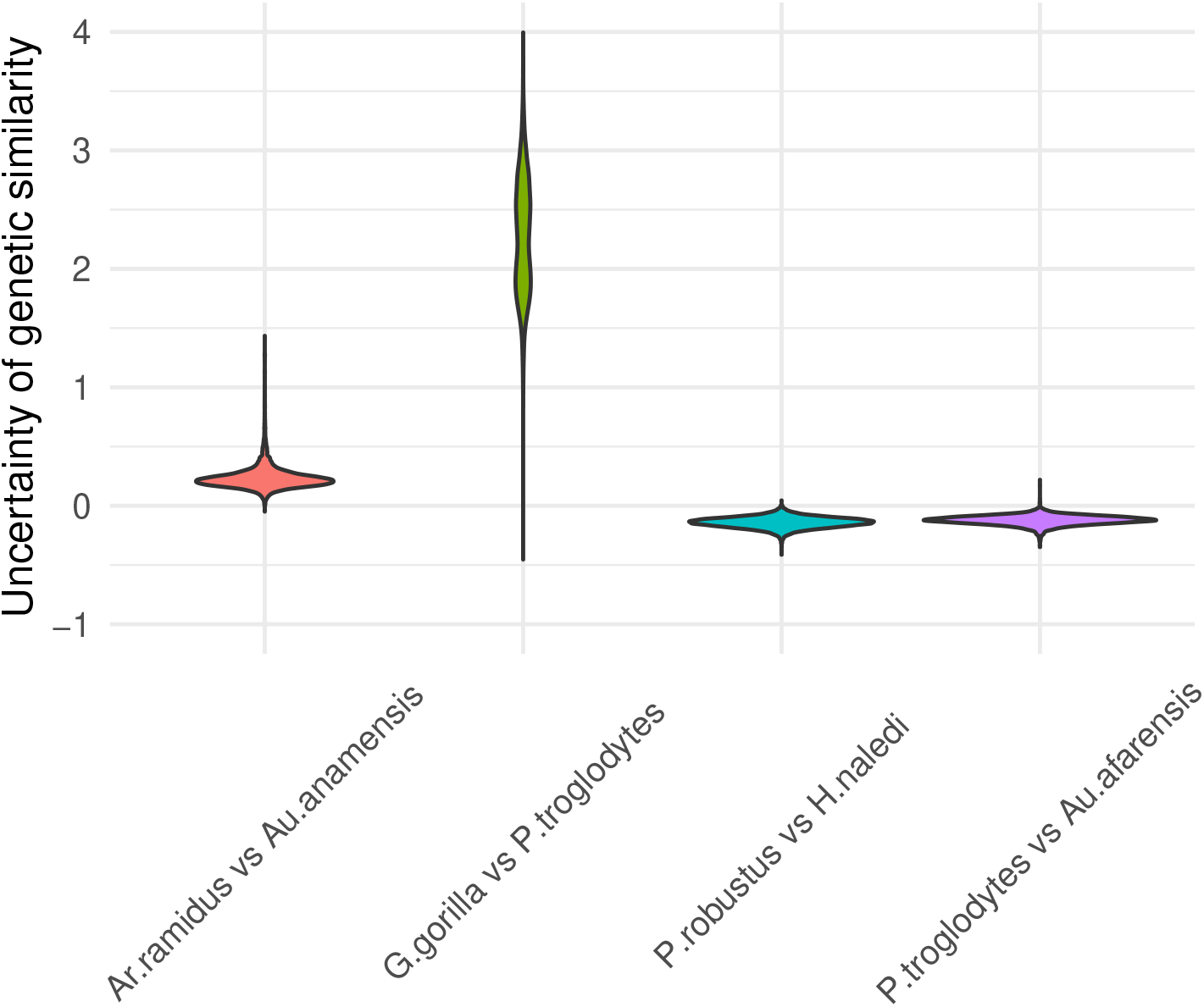
Uncertainty of genetic similarities between *Ar. ramidus* and *Au. anamensis, G. gorilla* and *P. troglodytes, P. robustus* and *H. naledi, P. troglodytes* and *Au. afarensis*.

## 4 Discussion

We provide an unbiased estimate for the genetic similarity matrix of samples, conditioned on a phylogenetic tree. This can be used to perform heritability and co-heritability estimates based on models such as Dahl et al. (2016), and can be used as a building block for LMM-like models in which uncertainty about inferred trees is modelled jointly with LMM regression parameters. As a proof-of-concept, we provide estimations of the genetic component for human and ancient hominin heights. To our knowledge, this is the first work to describe the integrals and expectations involved. The assumptions include haploidy (and no recombination), neutral models of evolution and the infinite sites model. These assumptions can be challenged by data in which samples undergo non-tree-like evolution, incomplete lineage sorting, hybridization or homoplasy. But these assumptions are appropriate for evolutionary time scales with limited lateral transfer (for example in hominins, where lateral transfer such as from Neanderthal to humans accounts for only a small proportion of the modern human genome; Sánchez-Quinto et al. 2012), or for trees derived from highly clonal bacteria such as *tuberculosis*.

Our numerical experiments demonstrate consistency between the empirically calculated genetic similarity matrix, and our proposed algorithm. We also apply our method to compute genetic blue for 8 hominin species, based on phylogenetic trees inferred from geological dates. Although our methods are based on the expected value of genetic similarity (*i.e*., molecular phylogeny), we assume that geological dates are proportional to molecular phylogeny. Our method can be used to provide genetic similarity matrices for heritability analyses (Dahl et al., 2016) in cases for which genotypes are not available. We also conduct a heritability analysis on the average heights of genetic males for hominin species. Even though the evolution of hominin species involves recombination and some lateral transfer, which is not accounted in our analysis, the main effects in molecular phylogeny at evolutionary timescales involve mutation at the species level.

In cases where genotypes are available, our work can be applied if phenotypes or geography suggests a mismatch between phylogeny and genotype. This can happen in application of linear mixed models for multivariate genomewide association studies with low-quality or low-coverage genotypes. This can also happen when genetic variation is low, but the examined phenotype is still under selection, causing homoplasy, which could bias estimates of a similarity matrix based on genotypes. When multiple samples of the phylogeny are available (for example, after running MCMC inference for the phylogeny), our algorithm can be used to compute the expected genetic similarity matrix for each posterior sample. The resulting matrices represents the uncertainty of genetic similarities among species, and could be combined with linear mixed models to better identify population stratification and correct for spurious association.

Our current approach is limited to the computation of genetic similarities among species, or samples without recombination. One future direction for this work would be to consider genealogies with recombination events. The ancestral recombination graph (ARG) describes the coalescence and recombination events among individuals (Rasmussen et al., 2014). The ARG is composed of a set of coalescent trees separated by break points. To compute expected genetic similarity matrix for samples given an ARG, we could first compute the expected genetic similarity matrices for each of the coalescent trees in the ARG, then compute weighted average for those expected genetic similarity matrices. The weights would be proportional to the number of loci between each consecutive pair of break points. This would constitute a straightforward extension of our method. Another future direction would be to incorporate more advanced evolutionary, mutation or selection models (relaxing the infinite sites and neutral assumptions). Finally, there are strong similarities between genetic and linguistic evolution (Cavalli-Sforza, 1997; Colonna et al., 2010). These two subjects both involve evolution and variation in a similar fashion, and phylogenetic methods have been applied to construct trees for language data (Atkinson and Gray, 2005; Jordan, 2007). Our work could be extended to compute similarities for languages using linguistic trees.

## Acknowledgements

We would like to thank Alexandre Bouchard Côté for helpful discussion. This research was supported by NSERC grant numbers RGPIN/05484-2019 and DGECR/00118-2019.

## Footnotes

1 https://github.com/shijiaw/Expected-Genetic-Similarity-Matrices

