## Supplementary Material for "Estimating Genetic Similarity Matrices using Phylogenies"

Shijia Wang[1] Shufei Ge[2] Caroline Colijn[3]
Priscila Biller[3] Liangliang Wang[4]
Lloyd T Elliott[4,*]

[1]School of Statistics and Data Science, LPMC and KLMDASR,
Nankai University, China

[2]Institute of Mathematical Sciences, ShanghaiTech University, China

[3]Department of Mathematics, Simon Fraser University, Canada

[4]Department of Statistics and Actuarial Science,
Simon Fraser University, Canada

### Appendix A

In this section, we describe an approach for estimating genetic similarities using phylogenies based on Gaussian kernels. Since branch lengths are proportional to the evolutionary events occurring along the branches, short branch length indicates close relatedness between a pair of samples. Given a phylogeny with $N$ samples, the genetic similarity between samples $i$ and $j$ is negatively proportional to the distance $d_{i,j}$ between them in the phylogeny. We define a Gaussian similarity by $K^S_{j,i} = \exp(-\lambda \cdot \frac{d_{i,j}}{|T|})$, where $|T|$ is the total branch length in a tree, and $\lambda$ is a bandwidth. This form is an adaption of the exponential isotropic model commonly used in spatial statistics, and yields positive definite $K^S$ (Gelfand et al., 2010). Algorithm 1 describes the computation of the Gaussian distance based genetic similarity.

**Algorithm** 1: Genetic similarity matrices from Gaussian distance

**Inputs:** A phylogenetic tree $t$ with a tree topology $\tau$ and a set of branch length $\boldsymbol{e} = (e_1, e_2, \ldots, e_{2N-3})$, and bandwidth $\lambda$.
**Output:** An $N \times N$ similarity matrix $K^S$.
Initialize $K^S_{ij} \leftarrow 0$.
Initialize pairwise distance and maximum pairwise distance with $d_{i,j} = 0$ and $|T| = 0$.
**for** $i \in \{1, 2, \ldots, N-1\}$ **do**
    **for** $j \in \{i+1, \ldots, N\}$ **do**
        Find the most recent common ancestor $c_{i,j}$ of $i$ and $j$ in $t$.
        Set $d_{i,j} \leftarrow l(c_{i,j}, i) + l(c_{i,j}, j)$. (Here $l(a, b)$ denotes the length of shortest path from $a$ to $b$ in $t$), and set $d_{j,i} \leftarrow d_{i,j}$.
        Set $|T| \leftarrow |T| + d_{i,j}$.
**for** $i \in \{1, 2, \ldots, N-1\}$ **do**
    Set $K_{i,j} = 1$.
    **for** $j \in \{i+1, \ldots, N\}$ **do**
        Set $K^S_{j,i} \leftarrow \exp(-\lambda \cdot d_{i,j}/|T|)$. Set $K^S_{j,i} \leftarrow K^S_{j,i}$.

### Appendix B

Figure S1 displays the entrywise difference between the expected genetic similarity matrices and the ground truth ($K^G_{ij} - K^T_{ij}$) with different number of taxa for each tree. The horizontal lines in a violin are 5%, 25%, 50%, 75%,

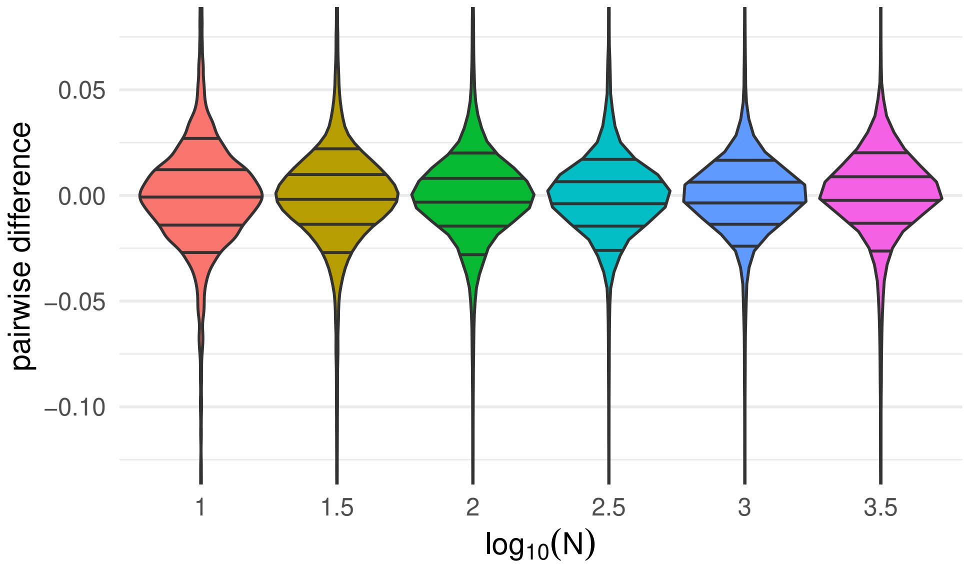


Figure S1: Comparison of simulated genetic similarity matrices from trees and genotypes as a function of number of taxa ($N$). The violins are provided for entrywise differences between the genetic similarity matrices produced by our method and the ground truth. The horizontal lines in a violin are 5%, 25%, 50%, 75%, and 95% quantiles respectively.

and 95% quantiles respectively. This figure indicates that the correlation among the entries of the genetic similarity matrices computed from trees and from genotypes is invariant to the number of taxa ($N$).

### Appendix C

Table 1 shows the expected genetic similarity matrix for the 8 hominin species considered in *Section 3.3* of the main text.

### Appendix D

In this section, we describe the details of the BEAST2 run for 24 hominin species (studied in *Section* 3.4). Time-calibrated phylogenies of 24 living and

Table 1: Genetic similarities for 8 hominin species.

| | *H. neanderthalensis* | *H. sapiens* | *H. antecessor* | *Asian H. erectus* | *African H. erectus* | *Georgian H. erectus* | *H. rudolfensis* | *H. habilis* |
|---|---|---|---|---|---|---|---|---|
| *H. neanderthalensis* | 0.493 | 0.400 | 0.077 | -0.116 | -0.153 | -0.187 | -0.233 | -0.283 |
| *H. sapiens* | 0.400 | 0.493 | 0.077 | -0.116 | -0.153 | -0.187 | -0.233 | -0.283 |
| *H. antecessor* | 0.077 | 0.077 | 0.756 | -0.104 | -0.141 | -0.175 | -0.221 | -0.271 |
| *Asian H. erectus* | -0.116 | -0.116 | -0.104 | 0.998 | -0.104 | -0.139 | -0.185 | -0.234 |
| *African H. erectus* | -0.153 | -0.153 | -0.141 | -0.104 | 1.078 | -0.129 | -0.175 | -0.224 |
| *Georgian H. erectus* | -0.187 | -0.187 | -0.175 | -0.139 | -0.129 | 1.176 | -0.156 | -0.206 |
| *H. rudolfensis* | -0.233 | -0.233 | -0.221 | -0.185 | -0.175 | -0.156 | 1.354 | -0.152 |
| *H. habilis* | -0.283 | -0.283 | -0.271 | -0.234 | -0.224 | -0.206 | -0.152 | 1.652 |

extinct hominids were inferred using BEAST v2.6.1 (Bouckaert et al., 2014). The dataset consists of 391 morphological characters partitioned according to their number of states (from Dembo et al. (2016)). Most characters (256) had binary states.

Each partition was assigned a site model with the appropriate Lewis MK substitution model (Lewis, 2001) with the same number of states as the number of states associated with characters in the partition. Each Lewis MK model had 4 Gamma categories, and the shape parameter of the Gamma distribution was estimated by BEAST.

The clock models and trees for each partition were linked (all partitions are considered to evolve on the same tree and under the same clock model). Branch lengths were allowed to vary under a relaxed clock exponential model. The rates associated with each branch were sampled from an exponential distribution whose parameters were also sampled by the MCMC. The tree prior was set to the fossilized birth-death (FBD) model, which estimates species divergence times from extant and fossil information in a coherent framework encompassing both diversification and fossil sampling.

The length of the MCMC was set to the default value (10,000,000), and 10,000 trees were uniformly subsampled from the posterior tree sample (thinning). Tracer v1.7.1 (Rambaut et al., 2018) was used to ensure that the posterior tree distribution has reached stationarity.

### Appendix E

Figure S2-S4 shows the posterior mean and the 95% credible interval of the genetic similarity matrix for the 24 hominin species considered in *Section 3.4*. Table 2 shows the index of hominin species in Figures S2-S4. The rows and columns of the heatmaps in Figures S2-S4 are displayed according to the following table.

Table 2: Index of hominin species in Figure S2-S4.

| | |
|---|---|
| 1 | G. gorilla, |
| 2 | P. troglodytes, |
| 3 | S. chadensis, |
| 4 | Ar. ramidus, |
| 5 | Au. anamensis, |
| 6 | Au. afarensis, |
| 7 | Au. africanus, |
| 8 | Au. garhi, |
| 9 | K. platyops, |
| 10 | P. robustus, |
| 11 | P. boisei, |
| 12 | P. aethiopicus, |
| 13 | Au. sediba, |
| 14 | H. habilis, |
| 15 | H. rudolfensis, |
| 16 | African H. erectus, |
| 17 | Asian H. erectus, |
| 18 | H. heidelbergensis, |
| 19 | H. neanderthalensis, |
| 20 | H. floresiensis, |
| 21 | H. sapiens, |
| 22 | H. antecessor, |
| 23 | Georgian H. erectus, |
| 24 | H. naledi |

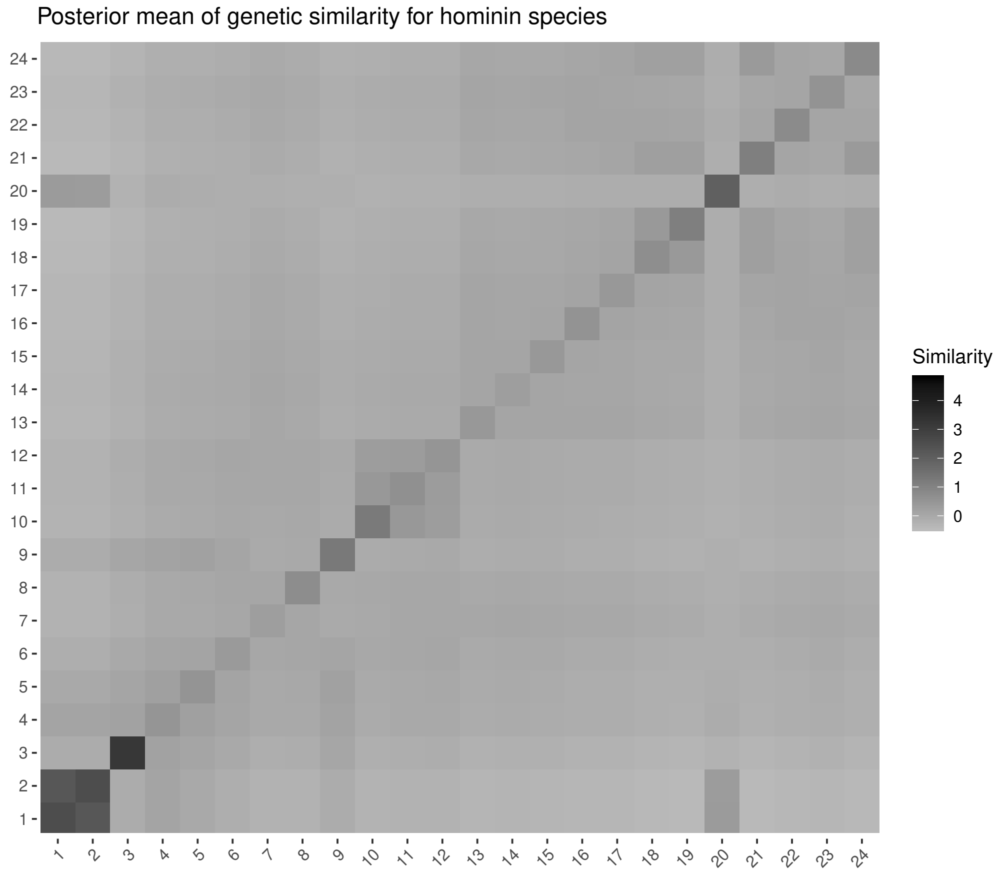


Figure S2: Posterior mean of genetic similarities for 24 hominin species.

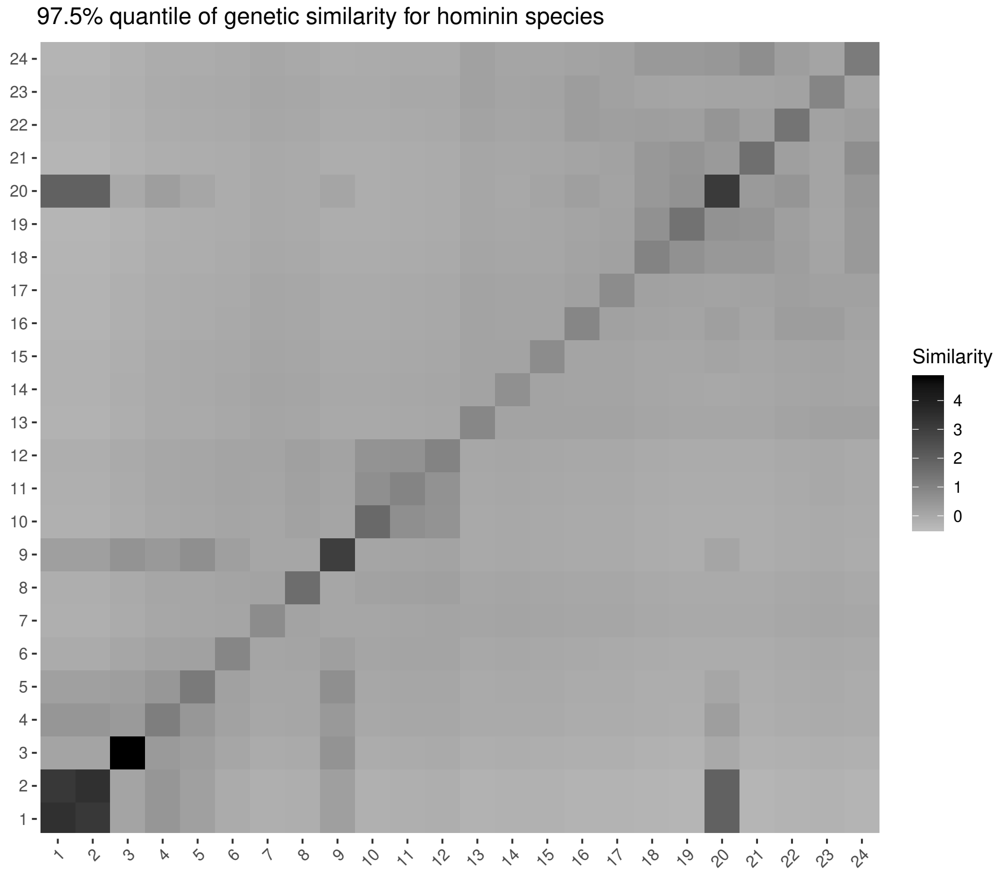


Figure S3: 97.5% quantile of genetic similarities for 24 hominin species.

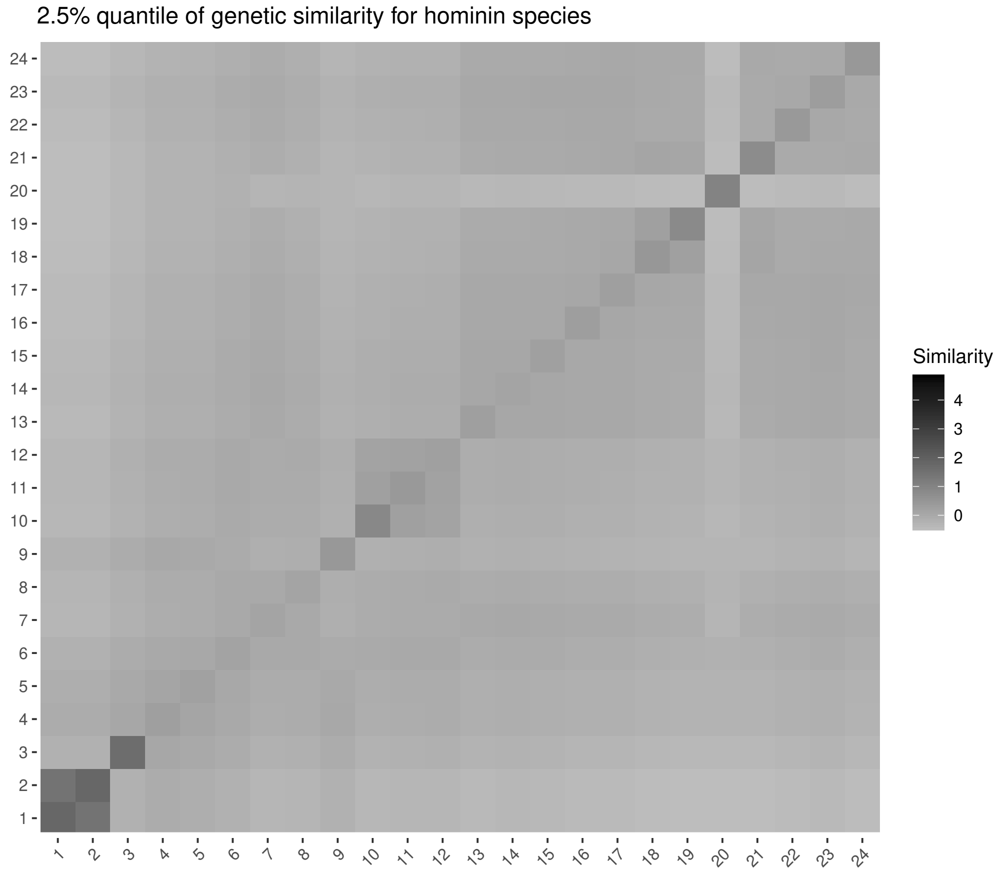


Figure S4: 2.5% quantile of posterior samples of genetic similarities for 24 hominin species.
